# STAT1 dissociates adipose tissue inflammation from insulin sensitivity in obesity

**DOI:** 10.1101/2020.04.10.036053

**Authors:** Aaron R. Cox, Natasha Chernis, David A. Bader, Pradip K Saha, Peter M. Masschelin, Jessica Felix, Zeqin Lian, Vasanta Putluri, Kimal Rajapakshe, Kang Ho Kim, Dennis T. Villareal, Reina Armamento-Villareal, Huaizhu Wu, Cristian Coarfa, Nagireddy Putluri, Sean M Hartig

**Affiliations:** Division of Diabetes, Endocrinology, and Metabolism, Baylor College of Medicine, Houston, TX; Department of Medicine, Baylor College of Medicine, Houston, TX; Department of Molecular and Cellular Biology, Baylor College of Medicine, Houston, TX; Dan L. Duncan Comprehensive Cancer Center, Advanced Technology Cores, Baylor College of Medicine, Houston, TX; Center for Translational Research on Inflammatory Diseases, Michael E DeBakey VA Medical Center, Houston, TX

## Abstract

Obesity fosters low-grade inflammation in white adipose tissue (WAT) that may contribute to the insulin resistance that characterizes type 2 diabetes mellitus (T2DM). However, the causal relationship of these events remains unclear. The established dominance of signal transducer and activator of transcription 1 (STAT1) function in the immune response suggests an obligate link between inflammation and the co-morbidities of obesity. To this end, we sought to determine how STAT1 activity in white adipocytes affects insulin sensitivity. STAT1 expression in WAT inversely correlated with fasting plasma glucose in both obese mice and humans. Metabolomic and gene expression profiling established STAT1 deletion in adipocytes (STAT1 fKO) enhanced mitochondrial function and accelerated TCA cycle flux coupled with subcutaneous WAT hyperplasia. STAT1 fKO reduced WAT inflammation, but insulin resistance persisted in obese mice. Rather, elimination of type I cytokine interferon gamma (IFN_γ_) activity enhanced insulin sensitivity in diet-induced obesity. Our findings reveal a permissive mechanism that bridges WAT inflammation to whole-body insulin sensitivity.

## INTRODUCTION

The obesity epidemic contributes to the increased health burden of chronic inflammatory conditions including insulin resistance, type 2 diabetes mellitus (T2DM), fatty liver, and cardiovascular disease ^1^. Obesity reflects facultative white adipose tissue (WAT) expansion that occurs during prolonged dietary stress. Although some clinical relationships explain how excess body weight causes insulin resistance in most individuals ^2^, WAT inflammation remains an enigmatic complication of obesity.

Obesity cultivates persistent low-grade inflammation that likely impacts the metabolic functions of WAT. Many studies demonstrate WAT inflammation causes local and systemic insulin resistance in rodents ^3–6^. In humans, expression of inflammatory cytokines in WAT durably correlate with body mass index (BMI) and insulin resistance ^7^. However, causal relationships between obesity-mediated inflammation and insulin resistance are still unclear, hindering the development of therapies that modulate the immune system to enhance the treatment of metabolic diseases arising concomitant with obesity.

In addition to mature adipocytes, WAT includes a dynamic stromal vascular fraction (SVF) containing lymphocytes and macrophages, which interact with one another and distant tissues through paracrine and endocrine signaling ^8^. Local elevation of the type I inflammatory cytokine interferon gamma (IFN_γ_) correlates with maladaptive WAT expansion and systemic insulin resistance ^9,10^. The IFN_γ_ receptor (IFN_γ_R1) communicates the IFN_γ_ signal through phosphorylation of Janus Kinases (JAKs) and subsequent activation of signal transducer and activator of transcription (STAT) proteins. STAT1 is the primary mediator of IFN_γ_ signaling and facilitates transcription of interferon-induced immune regulatory genes, including *IRF1*, *IRF9*, *ISG15* ^11–14^. Previously, we and others demonstrated inhibition of IFN_γ_ signaling in WAT prevents the development of insulin resistance and fatty liver in obese mice ^10,15–17^. Additional evidence suggests both type I and type II interferons signal to repress transcription factors essential for adipocyte differentiation and mitochondrial function ^18–21^. Therefore, while IFN_γ_ signaling plays an important role in the development of insulin resistance, the mechanistic underpinnings of this observation remain to be defined.

Here we used STAT1-deficient mouse models to investigate how the type I cytokine IFN_γ_ impacts insulin resistance and WAT inflammation. We found disruption of IFN_γ_-STAT1 signaling in mouse and human adipocytes relieves WAT inflammation and replenishes TCA metabolites. Mechanistically, STAT1 broadly represses gene programs involved in fatty acid metabolism and oxidative phosphorylation. As a result, STAT1 depletion in WAT increases subcutaneous adipocyte hyperplasia in obese mice. However, while adipocyte-specific STAT1 expression controls WAT inflammation, it is dispensable for systemic glucose control. Rather, complete deletion of IFN_γ_-STAT1 signaling using IFN_γ_gR1^−/-^ mice couples reduced inflammation with improved insulin sensitivity. We conclude STAT1 is a critical regulator of subcutaneous WAT expansion and the inflammatory response associated with obesity. These findings reveal how type I cytokine activity control obesity-associated inflammatory signals in adipose tissues.

## METHODS

### Animals

All procedures on animals have been approved by the Institutional Animal Care and Use Committee (IACUC) of Baylor College of Medicine (animal protocol AN-6411). All experiments were conducted using littermate-controlled male mice and were started when mice were aged 8-10 weeks. All experimental animals were maintained on a C57BL/6 background and housed in a barrier-specific pathogen-free animal facility with 12h dark-light cycle and free access to water and food. *Stat1^fl/fl^* mice were kindly provided by Lothar Henninghausen at the NIDDK ^22^. *Stat1^fl/fl^* mice were crossed with *AdipoQ-Cre* (Jackson Laboratory Stock No: 028020) to generate fat specific STAT1 knockout (*AdipoQ-Cre;Stat1^fl/fl^*, *STAT1*^fKO^) and littermate control (STAT^fl/fl^) mice. IFN_γ_gR1^−/-^ mice were purchased from Jackson Laboratory (Stock No: 003288) for breeding experimental cohorts in-house and C57BL/6J wild-type mice were obtained from the BCM Center for Comparative Medicine. Mice were fed 60% high fat diet (HFD; Research Diets) for 12 or 18 weeks before experiments.

### Human subjects

Subcutaneous WAT biopsies were obtained from 15 obese subjects during gastric bypass surgery ^15^. Nine subjects were defined as normal fasting plasma glucose (fasting plasma glucose=93.3±3.3 mg/dl, BMI=40.3±3.8, HOMA-IR=5.2±3.0), while 6 patients were defined as prediabetic (fasting plasma glucose=106.3±5.4 mg/dl, BMI=39.5±2.9, HOMA-IR=4.9±2.4). Additional subcutaneous WAT biopsies were obtained from 14 obese subjects ^23^. Nine subjects were defined as normal fasting plasma glucose (fasting plasma glucose=86.7±10.4 mg/dl), while 5 patients were insulin-resistant (fasting plasma glucose=110.8±4.1 mg/dl). Based on American Diabetes Association guidelines, normal fasting plasma glucose was defined as <100 mg/dl and prediabetes as fasting glucose of 100-125 mg/dl. One subject was excluded with a fasting plasma glucose of 152 mg/dl, identified as diabetic. Samples were stored at −80°C until RNA extraction. Human studies were approved by the Ethics Committee at Karolinska Institutet (Dnr 2008/2:3) and the New Mexico VA Health Care System (IRB: 11.139). All participants provided written informed consent.

### Mouse metabolic phenotyping

To determine glucose tolerance, mice were fasted for 16 hours and glucose was administered (1.5 g/kg body weight) by intraperitoneal (IP) injection. To determine insulin tolerance, mice were fasted four hours prior to insulin IP injection (1.5 U/kg body weight). Blood glucose was measured with a FreeStyle Freedom Lite Glucometer (Abbott Laboratories). Mouse body weight was measured weekly during HFD and body composition was examined by MRI (Echo Medical Systems). Overnight fasting serum levels were quantified by ELISA for insulin (#EZRMI-13K; Millipore) and leptin (#90030; Crystal Chem).

### Histology

Formalin-fixed paraffin-embedded adipose tissues sections were stained using anti-Mac3 (#550292; BD Biosciences) and hematoxylin & eosin (H&E) counterstain. Four 20x fields of view per tissue were imaged using a Nikon Ci-L Brightfield microscope and quantified using the Adiposoft ImageJ plugin ^24^ for adipocyte cell size.

### Antibodies and Immunoblotting

Cell and tissue lysates were prepared in Protein Extraction Reagent (Thermo Fisher) supplemented with Halt Protease and Phosphatase Inhibitor Cocktail (Thermo Fisher). Western blotting was performed with whole cell lysates run on 4-12% Bis-Tris NuPage gels (Life Technologies) and transferred onto Immobilon-P Transfer Membranes (Millipore) followed by antibody incubation (antibodies listed in **Supplementary Table 1)**. Immunoreactive bands were visualized by chemiluminescence.

### Real-time qPCR

Total RNA was extracted using the Direct-zol RNA MiniPrep kit (Zymo Research). cDNA was synthesized using SuperScript VILO Master Mix (Thermo Fisher). Relative mRNA expression was measured with Taqman Fast qPCR reagents using a QuantStudio 3 real-time PCR system (Applied Biosystems). Invariant controls included *TBP*. TaqMan and Roche Universal Probe Gene Expression Assays are detailed in **Supplementary Table 2**.

### RNA-seq

Sample quality control, mRNA library prep, and RNA sequencing was performed by the University of Houston Sequencing and Editing Core. RNA sample quality control assessment (RNA integrity number ≥8) was performed with the RNA Pico 6000 chip on Bioanalyzer 2100 (Agilent) and were quantified with Qubit Fluorometer (Thermo Fisher). mRNA libraries were prepared with Ovation RNA-Seq System V2 (NuGen) and Ovation Ultralow Library System V2 (NuGen) using input RNA. Size selection for libraries was performed using SPRIselect beads (Beckman Coulter) and purities of the libraries were analyzed using the High Sensitivity DNA chip on Bioanalyzer 2100 (Agilent). The prepared libraries were pooled and sequenced using Illumina NextSeq 500 generating ~20 million 76bp paired-end reads/sample. Reads were mapped to the UCSC mouse reference genome mm10 using HISAT2 ^25,26^. Stringtie was used to calculate the expression level as reads per kilobase per million (RPKM). Gene set enrichment analysis was performed with Molecular Signatures Database and normalized enrichment scores were calculated for the Hallmark gene sets. Hierarchical cluster analysis was performed by Euclidean distance using Log_2_ fold change over STAT1^fl/fl^ controls of significantly altered (p<0.05) genes. RNA-seq data set can be accessed through Gene Expression Omnibus.

### Metabolomics

For extraction of inguinal WAT metabolites, 750 μl of water/methanol (1:4) was added to snap-frozen WAT and samples were homogenized, then mixed with 450 μl ice-cold chloroform. The resulting solution was mixed with 150 μl ice-cold water and vortexed again for 2 minutes. The solution was incubated at –20°C for 20 minutes and centrifuged at 4°C for 10 minutes to partition the aqueous and organic layers. The aqueous and organic layers were combined and dried at 37°C for 45 minutes in an automatic Environmental Speed-Vac system (Thermo Fisher Scientific). The extract was reconstituted in a 500-μl solution of ice-cold methanol/water (1:1)extract was resuspended in and filtered through a 3-kDa molecular filter (Amicon Ultracel 3-kDa Membrane) at 4°C for 90 minutes to remove proteins. The filtrate was dried at 37°C for 45 minutes in a speed vacuum and stored at – 80°C until MS analysis. Prior to MS analysis, the dried extract was resuspended in a 50-μl solution of methanol/water (1:1) containing 0.1% formic acid and then analyzed using multiple reaction monitoring (MRM). Ten microliters were injected and analyzed using a 6490 QQQ triple quadrupole mass spectrometer (Agilent Technologies) coupled to a 1290 Series HPLC system via selected reaction monitoring (SRM).

Separation of glycolytic and TCA intermediates. Briefly, aqueous phase chromatographic separation was achieved using three solvents: water (solvent A), water with 5mM ammonium acetate (pH 9.9), and 100% acetonitrile (ACN) (solvent B). The binary pump flow rate was 0.2 ml/min with a gradient spanning 80% B to 2% B over a 20-minute period followed by 2% B to 80% B for a 5 min period and followed by 80% B for 13-minute time period. The flow rate was gradually increased during the separation from 0.2 mL/min (0-20 mins), 0.3 mL/min (20-25 min), 0.35 mL/min (25-30 min), 0.4 mL/min (30-37.99 min), and finally set at 0.2 mL/min (5 min). Glycolytic and TCA intermediates were separated on a Luna Amino (NH2) column (3 µm, 100A 2 × 150 mm, Phenomenex), that was maintained in a temperature-controlled chamber (37°C).

Glycolytic and TCA intermediates were measured using negative ionization mode with an ESI voltage of −3500ev. Approximately 9–12 data points were acquired per detected metabolite. For all samples, ten microliters of sample were injected and analyzed using a 6495 QQQ triple quadrupole mass spectrometer (Agilent) coupled to a 1290 series HPLC system via SRM. The data was normalized with internal standard, log2-transformed, and per-sample basis. For every metabolite in the normalized dataset, t-tests were conducted to compare expression levels between different groups. Differential metabolites were identified by adjusting the p-values for multiple testing at an FDR threshold of <0.25.

### In vitro experiments

Subcutaneous primary human preadipocytes were provided by Zen-Bio, differentiated, and transfected as previously described ^27^. Mature adipocytes were transfected with STAT1 siRNA (GE Dharmacon), siRNA control (GE Dharmacon) at a final concentration of 50 nM. Recombinant human and mouse IFN_γ_ were purchased from R&D Systems. FLAG-*STAT1* (Addgene #71454) or vector control lentiviral particles were prepared and introduced into human adipocytes for 48h before experiments.

For CRISPR-Cas9 gene deletion experiments, single guide RNAs (gRNA) targeting sequences in exon 7 (g1), exon 6 (g2), and exon 17 (g3) of *Stat1* were designed using the Broad Institute GPP Web Portal. A non-mammalian targeting control sgRNA sequence with similar GC content was used as a control ^28^. Guide sequences (listed in **Supplementary Table 3**) were cloned into the lentiCRISPR v2 plasmid (Addgene #52961) and lentiviral particles were generated in 293T cells (ATCC) using packaging plasmids pMD2.G (Addgene #12259) and psPAX2 (Addgene #12260) for transfection with iMFectin (Gendepot). 3T3-L1 cells were selected for vector incorporation using puromycin (Gibco). Cas9 expression and STAT1 disruption were confirmed via qPCR and Western blotting with g3 inducing complete loss of STAT1.

### Adipocytes differentiated from stromal vascular fractions

Stromal vascular fractions (SVFs) were isolated from mouse inguinal WAT. Fat depots were digested in PBS containing collagenase D (Roche, 1.5 U/ml) and dispase II (Sigma, 2.4 U/ml) supplemented with 10 mM CaCl_2_ at 37°C for 40-45 min. The primary cells were filtered twice through 70 μm cell strainers and centrifuged at 700 rcf to collect the SVF. The SVF cell pellets were rinsed and plated. Adipocyte differentiation was induced by treating confluent cells in DMEM/F12 medium containing glutamax (ThermoFisher), 10% FBS, 0.250 mM isobutylmethylxanthine (Sigma Chemical Co.), 1 μM rosiglitazone (Cayman Chemical Co.), 1 μM dexamethasone (Sigma Chemical Co.), 850 nM insulin (Sigma Chemical Co.), and 1 nM T3 (Sigma Chemical Co). Four days after induction, cells were switched to the maintenance medium containing 10% FBS, 1 μM rosiglitazone, 1 μM dexamethasone, 850 nM insulin, 1 nM T3. Experiments that tested IFN_γ_ effects on SVF-derived adipocytes occurred 8-10 days after induction of differentiation.

### Cellular Respiration

Respiration was measured in adipocytes using an XF24 analyzer (Seahorse Bioscience). Preadipocytes were plated into V7-PS plates and differentiated before treatments. For the assay, media was replaced with 37°C unbuffered DMEM containing 4.5 g/L glucose, sodium pyruvate (1 mmol/L), and L-glutamine (2 mmol/L). Basal respiration was defined before sequential addition of forskolin, oligomycin, rotenone, and antimycin A.

### Statistical Analyses

Statistical significance was assessed by unpaired one-sided Student’s t-test or Mann-Whitney U test. All data are presented as mean SEM. All tests were carried out at the 95% confidence interval, unless otherwise stated. Pearson’s correlation coefficient (Pearson’s r) was calculated to evaluate correlations between metabolic parameters and *STAT1* expression in human subcutaneous WAT.

### Data and Resource Availability

The datasets and resources generated during the current study are available from the corresponding author upon reasonable request. RNA-seq data deposited in Gene Expression Omnibus (accession number: pending).

## RESULTS

### STAT1 expression corresponds with impaired glucose and fat metabolism in human and mouse adipocytes

STAT1 regulates genes that drive pro-inflammatory responses ^11–14^ and impair mitochondrial function ^15,29^. To test the hypothesis that *STAT1* expression in WAT inversely correlates with insulin sensitivity, we measured *STAT1* levels in fat tissues collected from obese mice and humans. *STAT1* expression was higher in subcutaneous WAT isolated from diet-induced obesity (DIO) mice compared to mice fed normal chow (Figure 1A). Similarly, in obese humans, *STAT1* was almost three-fold higher (Mann-Whitney U, p=0.092) in subcutaneous WAT isolated from subjects with prediabetes (fasting plasma glucose 100-125 mg/dl) compared to normoglycemic (fasting plasma glucose <100 mg/dl) counterparts (Figure 1B). Likewise, *STAT1* levels correlated with fasting plasma glucose (Pr=0.322, p=0.088). These results indicate increased *STAT1* in WAT expression is associated with insulin resistance.

**Figure 1.**
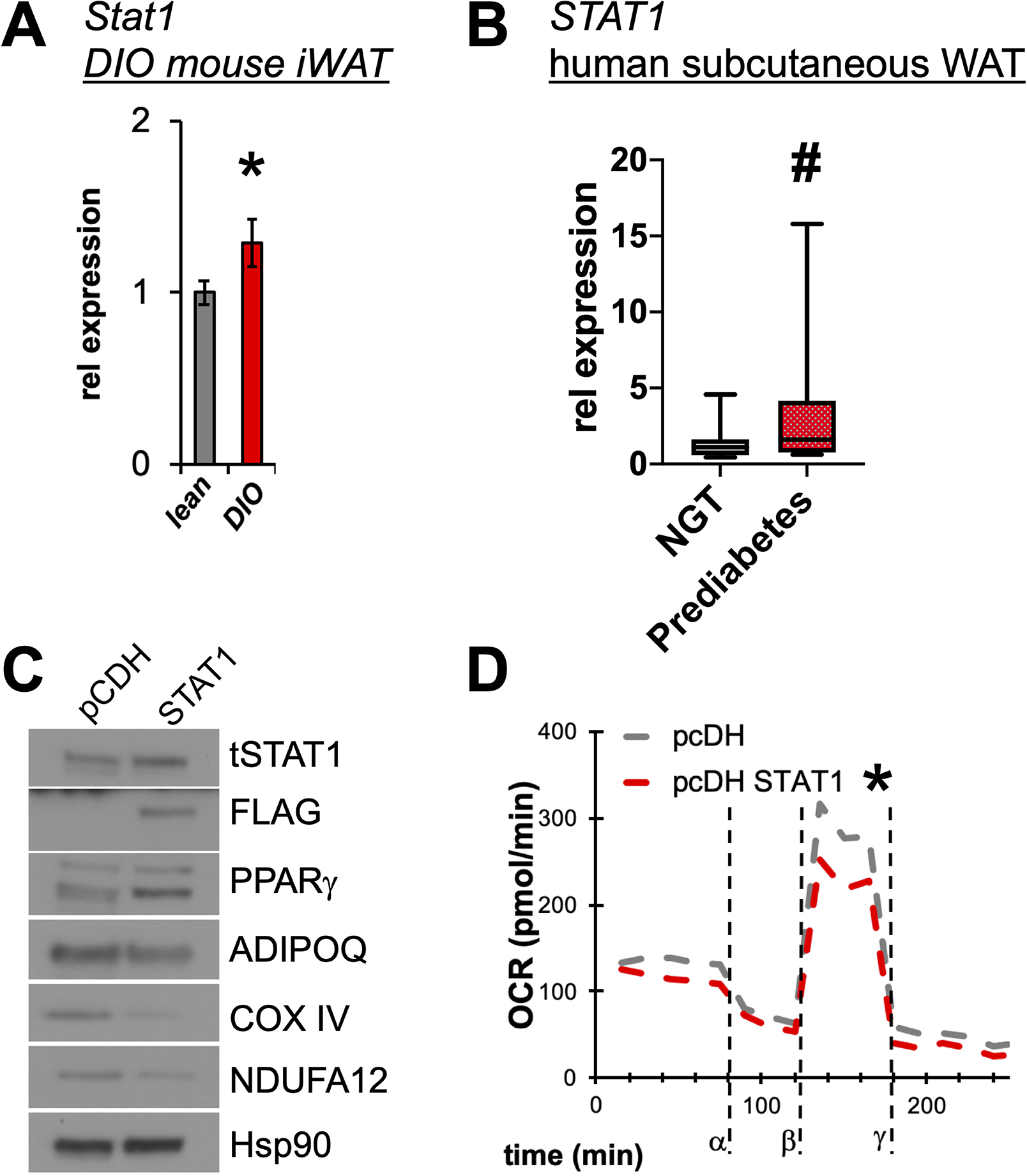
Higher STAT1 levels correspond with impaired adipocyte lipid metabolism. **(A)** Relative mRNA expression of *Stat1* in lean (grey) or high fat diet-induced obese (DIO) (red) wild-type mice (n=4/group). *p<0.05, data are represented as mean +/− SEM. **(B)** Relative *STAT1* expression was measured in human subcutaneous adipose tissue biopsied from subjects with prediabetes (red; n=11) compared to those with normal glucose tolerance (NGT, grey; n=18) #p<0.09). Human subcutaneous preadipocytes were differentiated for 8 days and then transfected with *STAT1* or control vector (pcDH). *p<0.05, data are represented as mean +/− SEM. **(C)** To confirm STAT1 expression, immunoblotting of total STAT1 and FLAG was performed, along with markers of mature adipocytes and mitochondrial proteins. **(D)** Respiration (as oxygen consumption rate, OCR) was measured in human adipocytes expressing control vector (pcDH; grey line) or *STAT1* (red line) over time with the addition of oligomycin (α), carbonyl cyanide-*4*-(trifluoromethoxy) phenylhydrazone (FCCP) (β), and antimycin-A/rotenone (γ) (n=5). *p<0.05 indicates changes in maximal respiration.

To establish STAT1 as a negative regulator of metabolic functions, we overexpressed FLAG-tagged STAT1 in primary human adipocytes. Enforced STAT1 expression repressed the level of adiponectin (ADIPOQ), a surrogate for insulin sensitivity. Adipocyte metabolism and insulin sensitivity correlate with expression of mitochondrial electron transport chain (ETC) components ^30^. Accordingly, STAT1 repressed mitochondrial ETC proteins (COXIV and NDUFA12), suggesting impaired respiratory capacity (Figure 1C). We next measured oxygen consumption rate (OCR) by Seahorse to establish the functional impact of these changes on metabolic activity. We found STAT1 overexpression suppressed maximal respiration in human adipocytes compared to control infections (Figure 1D). Together, these results demonstrate higher STAT1 expression impairs metabolic functions of adipocytes.

### STAT1 depletion improves human and mouse fat cell function

To further define the metabolic impact of STAT1 on adipocyte function, we leveraged gene editing and siRNA approaches to deplete STAT1 in primary human and 3T3-L1 mouse adipocytes. First, we transfected mature human adipocytes with scrambled control (scRNA) or *STAT1* siRNA followed by treatment with vehicle or IFN_γ_ for 24 h (Figure 2A). *STAT1* knockdown blocked IFN_γ_-mediated induction of *STAT1* itself, as well as the *STAT1* target gene *IRF1*. Remarkably, depletion of *STAT1* increased basal expression of genes that reflect insulin sensitivity (*ADIPOQ, UCP1*) and mediated resistance to the effects of IFN_γ_ treatment. Likewise, IFN_γ_ treatment decreased maximal adipocyte OCR in control cells (green lines), but STAT1 knockdown (red lines) conferred resistance to this response (Figure 2B).

**Figure 2.**
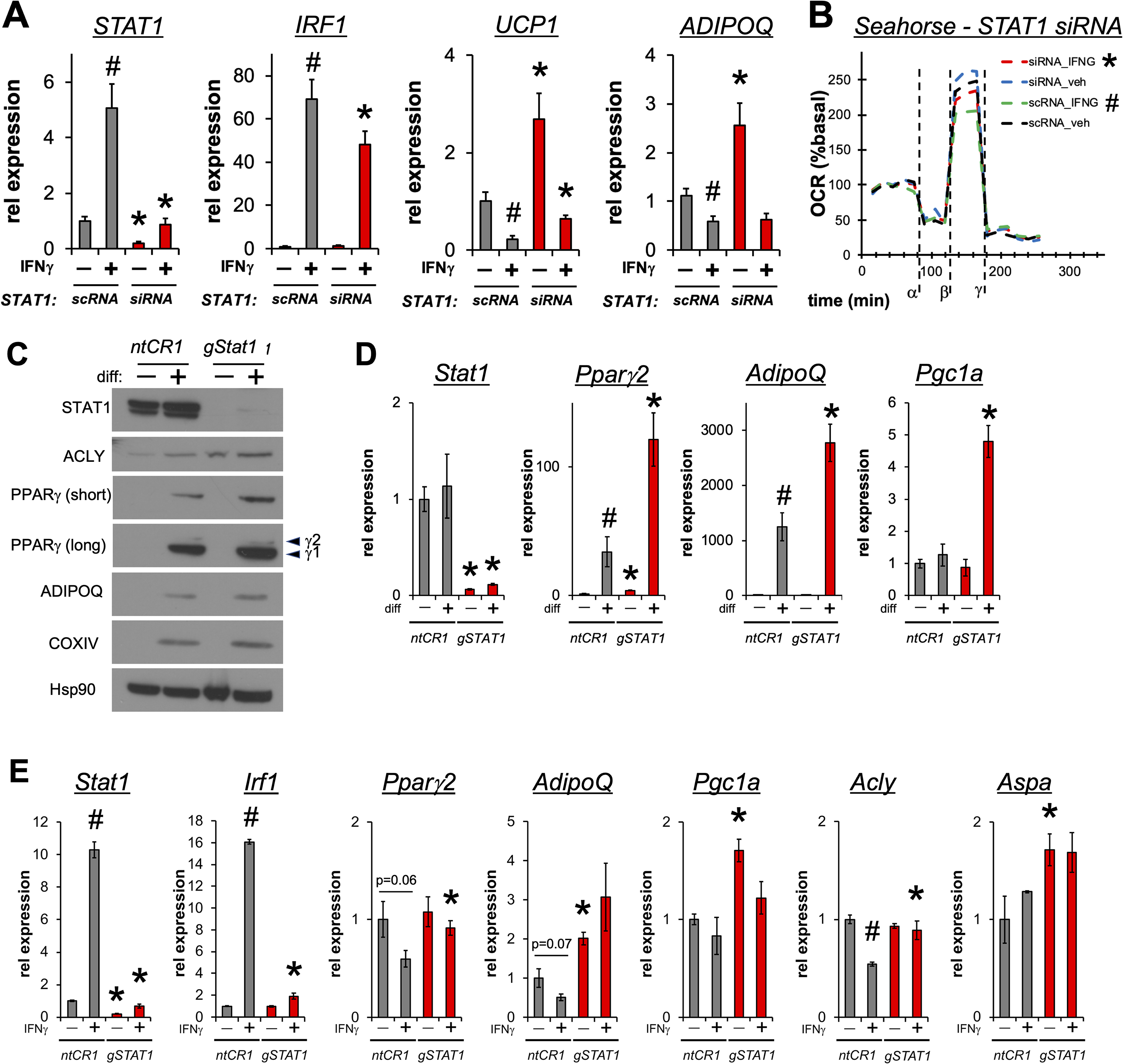
STAT1 deletion restores adipocyte metabolism in human and mouse adipocytes. Human subcutaneous preadipocytes were differentiated for 10 days and then transfected with *STAT1* siRNA or scramble RNA (scRNA; control) for 48h. After transfection, cells were treated +/− 100 ng/ml IFN_γ_ for 24h. **(A)** Relative mRNA expression of *STAT1*, *IRF1*, *UCP1*, and *ADIPOQ* (n=9). grey – scRNA, red – STAT1 siRNA; *p<0.05 vs scRNA, #p<0.05 vs -IFN_γ_ vehicle treated; data are represented as mean +/− SEM. **(B)** Oxygen consumption rate (OCR) in differentiated human adipocytes after exposure to IFN_γ_ with addition of oligomycin (α), carbonyl cyanide-*4*-(trifluoromethoxy) phenylhydrazone (FCCP) (β), and antimycin-A/rotenone (γ). (black – scRNA vehicle, green – scRNA +IFN_γ_, blue – *STAT1* siRNA vehicle, red – *STAT1* siRNA + IFN_γ_) (n=12); *p<0.06 vs scRNA, #p<0.06 vs -IFN_γ_ vehicle treated for maximal respiration. 3T3-L1 cells were transfected with Cas9 and one of three *Stat1* single guide RNAs (g1, g2, g3) or a non-mammalian targeting control (ntCR1) guide RNA. **(C)** Immunoblots of total lysates from ntCR1 or g3 *Stat1* (gSTAT1) cells +/− differentiation for 10 days. **(D)** Relative mRNA expression of *Stat1*, *Ppar_γ_2*, *AdipoQ*, and *Pgc1a* from ntCR1 and g3 *Stat1* cells +/− differentiation for 10 days (n=3). **(E)** Differentiated ntCR1 and gSTAT1 cells were treated +/− 100 ng/ml IFN_γ_ for 24h and then harvested for quantification of relative mRNA for inflammatory (*Stat1*, *Irf1*), fat cell identity (*Ppar_γ_2*, *AdipoQ*), and lipid metabolism genes (*Pgc1a*, *Acly*, *Aspa*) (n=3). grey – ntCR1; red – gSTAT1; *p<0.05 vs ntCR1, #p<0.05 vs vehicle; data are represented as mean +/− SEM.

To examine the impact of *STAT1* signaling during adipocyte differentiation, we leveraged CRISPR-Cas9 to knock out *STAT1* in 3T3L1 cells. We screened three single guide RNAs in 3T3-L1 cells targeting different exons of *Stat1* and determined *Stat1* guide #3 (g3) targeting exon 17 imposed complete depletion of Stat1 protein expression. *Stat1* knockout significantly enhanced induction of 3T3-L1 adipocyte maturation as indicated by elevated expression of proteins classically associated with adipocyte differentiation (PPAR_γ_, ADIPOQ, ACLY). Similarly, COXIV, a subunit of the terminal enzyme of the mitochondrial ETC, was increased in *Stat1* knockout cells, suggesting enhanced respiratory capacity in these cells (Figure 2C). *Stat1* knockout also increased the expression of PPAR_γ_ target genes including *Ppar_γ_2*, *AdipoQ*, and *Pgc1a* (Figure 2D). To test how *Stat1* knockout affects the response to inflammatory stimulus, we treated control (ntCR1) and *STAT1* knockout (gSTAT1) adipocytes with IFN_γ_ for 24 h (Figure 2E). IFN_γ_ treatment stimulated expression of *Stat1* and *Irf1* and suppressed the adipocyte marker genes *Ppar_γ_*2, *AdipoQ*, and *Acly* in control cells. In contrast, *Stat1* deletion increased expression of *AdipoQ*, *Pgc1a*, and the lipogenic enzyme *Aspa*, and reversed IFN_γ_ repression of *Ppar_γ_*2 and *Acly*. Collectively, these experiments performed in human and mouse adipocytes demonstrate STAT1 inhibition elevates mitochondrial function, accelerates adipocyte differentiation, and blunts responses to the obesity-related cytokine IFN_γ_.

### STAT1 deletion in adipocytes induces subcutaneous fat cell hyperplasia in mice

STAT1 regulates inflammatory responses in multiple tissues and cell types ^31–34^. However, the immune cell functions of STAT1 in obesity may mask dominant roles in WAT. Therefore, to examine the adipocyte specific impact of STAT1 signaling on obesity-induced inflammation and insulin resistance, we crossed *STAT1^fl/fl^* mice ^22^ with mice expressing Cre recombinase under control of the adiponectin promoter ^35^ to generate STAT1 fat-specific knockout mice (*STAT1*^fKO^). Immunoblotting confirmed tissue-specific loss of STAT1 expression in WAT depots from *STAT1*^fKO^ mice (Figure 3A). Liver STAT1 levels and expression of STAT2/3 were unchanged in *STAT1*^fKO^ mice, validating specific deletion of STAT1 only in tissues that express adiponectin (Figure 3B). mRNA analysis confirmed adipocyte-specific knockout reduced *Stat1* levels by 70% in WAT (Figure 3C). To model obesity and the metabolic stress resulting from excessive caloric intake, *STAT1*^fKO^ mice and littermate controls were maintained on a HFD for 18 weeks. Contrary to our expectations, weight gain (Figure 3D) and body composition (Figure 3E) were similar between groups after diet-induced obesity. We observed nominal improvements in insulin (Figure 3F) and glucose tolerance (Figure 3G), as well as overnight fasted serum insulin levels (Figure 3H) in *STAT1*^fKO^ mice compared to controls.

**Figure 3.**
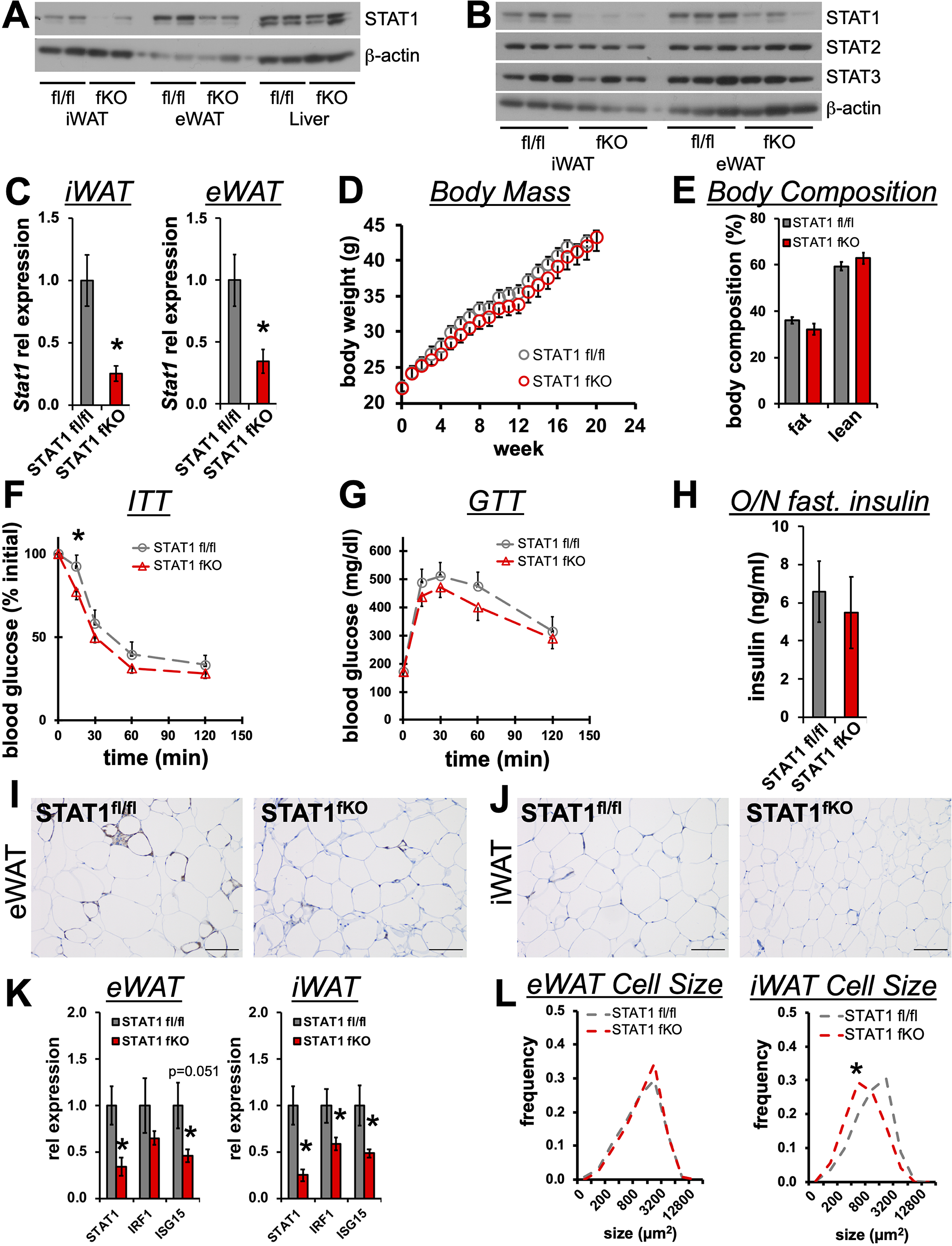
STAT1 deletion in adipocytes induces subcutaneous fat cell hyperplasia. **(A)** Immunoblots show STAT1 knockdown in iWAT and eWAT, but not liver, of fat specific (ADIPOQ-Cre) *STAT1^fKO^* compared to *STAT1^fl/fl^* littermate controls. **(B)** Immunoblots show unaltered STAT2 and STAT3 expression in iWAT and eWAT of *STAT1^fKO^* mice. **(C)** Relative *Stat1* expression in the iWAT and eWAT of *STAT1^fl/fl^* and *STAT1^fKO^* mice (n=7-8/group). **(D)** Body mass and **(E)** composition (% body mass) for *STAT1^fl/fl^* and *STAT1^fKO^* mice on HFD for 18 weeks (n=9-11/group). **(F-H)** Insulin (ITT; n=12-13/group) and glucose (GTT; n=8/group) tolerance tests with corresponding overnight fasting serum insulin (n=8/group) in *STAT1^fl/fl^* and *STAT1^fKO^* mice on HFD. **(I)** eWAT and **(J)** iWAT staining for macrophages (Mac3; brown) and **(K)** relative mRNA expression of inflammatory genes. **(L)** Adipocyte cell size distribution (% total cells) tabulated across four 20x fields of view per mouse fat depot (n=5-6/group) grey – *STAT1^fl/fl^*; red – *STAT1^fKO^* *p<0.05, data are represented as mean +/− SEM.

Diet-induced obesity increases macrophage infiltration of epididymal (eWAT) and inguinal (iWAT) adipose depots associated with impaired WAT hyperplasia ^36^. Based on our *in vitro* studies, adipocyte-specific deletion of *STAT1* may decrease the production of pro-inflammatory signaling molecules in WAT, thereby restoring adipocyte differentiation capacity. To examine this hypothesis, we collected the iWAT and eWAT depots from the *STAT1*^fKO^ mice and littermate controls for immunohistochemistry and molecular analysis. Grossly, STAT1 deletion in the adipocytes of obese mice decreased pro-inflammatory macrophage infiltration into eWAT (Figure 3I) and iWAT (Figure 3J) while reducing mRNA expression of several inflammatory STAT1 target genes in *STAT1^fKO^* WAT compared to controls (Figure 3K). Last, we performed quantitative image-based histological analysis to compare the average adipocyte size in STAT1^fKO^ WAT compared to controls. We observed adipocytes from *STAT1*^fKO^ subcutaneous iWAT were significantly smaller, favoring healthy expansion of this depot through adipocyte hyperplasia rather than hypertrophy (Figure 3L). Together, these results suggest STAT1 disruption in adipocytes reduces inflammation in WAT, which facilitates healthy WAT expansion and may in turn improve adipocyte metabolism.

### Subcutaneous WAT from *STAT1^fKO^* mice exhibit enrichment of mitochondrial genes and TCA cycle improvements

Our mouse and in vitro models suggest STAT1-deficient fat cells engage pathways that merge metabolic and differentiation genes to enable hyperplasia in the setting of obesity. Therefore, we used RNA-Seq to identify the biologically cohesive gene programs of STAT1 depletion in the WAT of obese *STAT1^fl/fl^* and *STAT1^fKO^* mice. These efforts uncovered clear signatures that explain the impacts of STAT1 knockout in the eWAT and iWAT (Figure 4A). In *STAT1^fKO^* mice, iWAT and eWAT shared 22 suppressed genes, 11 of which were common IFN_γ_-STAT1 targets (e.g. *Stat1*, *Stat2*, *Isg15*). Consistent with its known roles in inflammation, Gene Set Enrichment Analysis (GSEA) indicated *STAT1* deletion in WAT exerted broad anti-inflammatory effects including evidence for suppression of interferon responses (Figure 4B). Expression of STAT1 targets within these genes sets (*Stat1*, *Irf1*, *Isg15*, *Oas1*) was reduced in *STAT1^fKO^* iWAT (Figure 4C). Consistent with the idea that reduced inflammation may improve adipocyte function, GSEA also revealed *STAT1^fKO^* increased levels of genes found in central metabolic pathways including oxidative phosphorylation, adipogenesis, and fatty acid metabolism. Notably, a distinct series of lipid metabolism and mitochondrial genes were selectively enhanced in the iWAT of *STAT1^fKO^* compared to controls (Figure 4D). Together, these findings explain how STAT1 deletion imparts gene changes that enhance adipocyte metabolism and hyperplasia.

**Figure 4.**
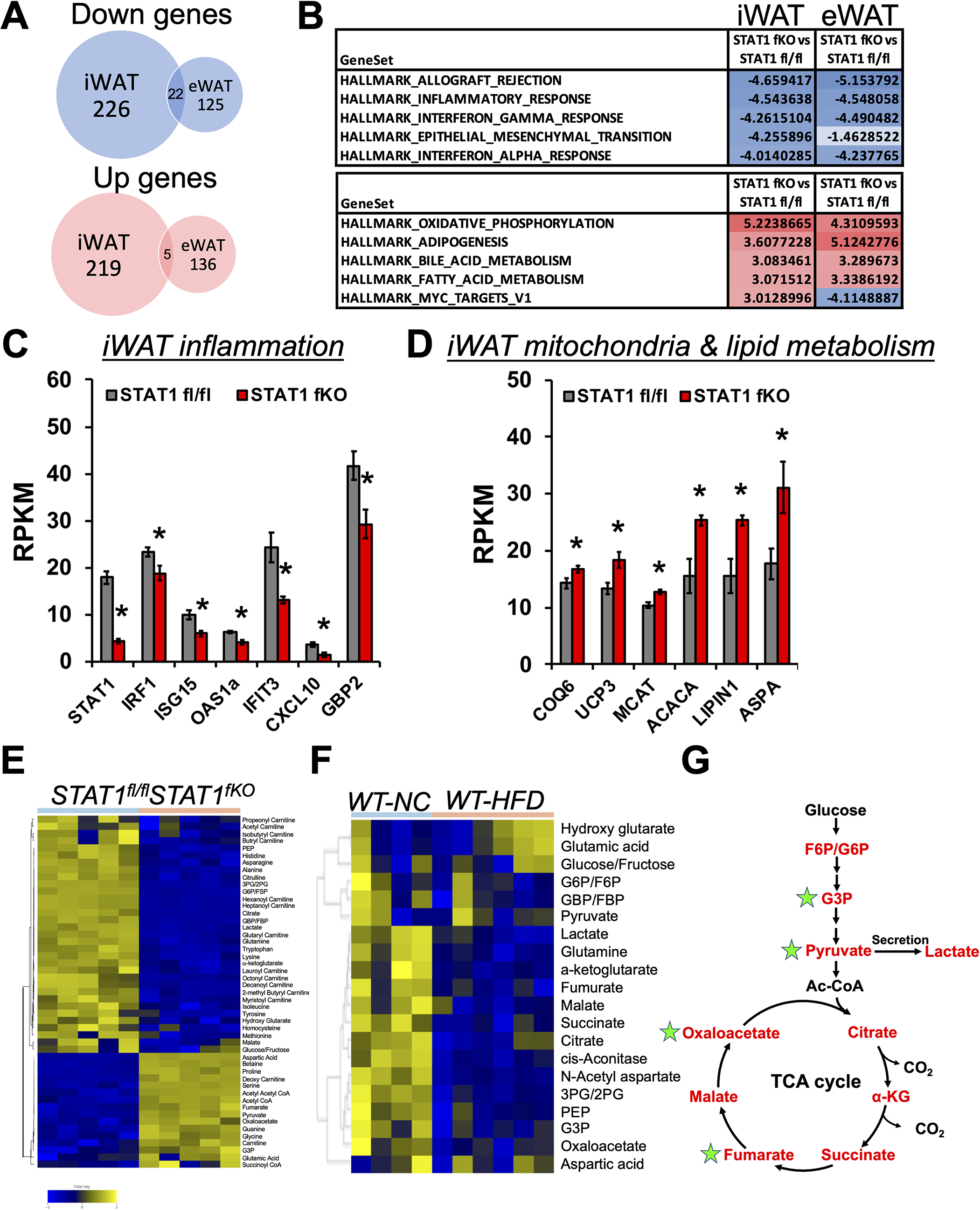
Inguinal WAT from STAT1 fKO mice exhibit enrichment of mitochondrial genes and TCA cycle improvements. **(A)** RNA-Seq coupled with **(B)** GSEA identified gene signatures altered by STAT1 knockout in the eWAT and iWAT of obese mice. Relative mRNA expression of key genes that validate the **(C)** anti-inflammatory and **(D)** metabolic gene signatures of STAT1 deletion in iWAT. **(E)** Heatmap representing hierarchical clustering of differential metabolites in iWAT between obese *STAT1^fl/fl^* and *STAT1^fKO^* mouse models (n=5/group; FDR<0.25) and **(F)** lean or diet-induce obese wild-type mice (n=4-6/group) were assessed using mass spectrometry. **(G)** Metabolomics analysis of iWAT establish HFD feeding in WT mice reduced (red) tricarboxylic acid (TCA) cycle metabolites that become rescued (green) in *STAT1^fKO^* mice. G6P/F6P – glucose/fructose-6-phosphate; GBP/FBP – glucose/fructose-1,6-bisphosphate; 3PG/2PG – 3-/2-phosphoglycerate; G3P – glyceraldehyde 3-phosphate; PEP – phosphoenolpyruvate; Ac-CoA – acetyl-CoA; -KG – alpha-ketoglutarate. grey – *STAT1^fl/fl^*; red – *STAT1^fKO^*; *p<0.05, data are represented as mean +/− SEM.

To further understand the metabolic impact of *STAT1* deletion in obesity, we assessed the relative steady state levels of metabolic intermediates in iWAT collected from obese *STAT1^fl/fl^* and *STAT1^fKO^* mice after 18 weeks HFD (Figure 4E). To identify specific metabolic pathways targeted by STAT1 in adipocytes, we performed a broad analysis of glycolytic, TCA cycle, acylcarnitines, and amino acid metabolites. We observed depletion of several long chain fatty acid carnitine species (myristoyl carnitine, octenyl carnitine, lauroyl carnitine) from iWAT of *STAT1^fKO^* mice that reflect higher rates of beta oxidation 37. Moreover, *STAT1* loss caused accumulation of pyruvate and key metabolic intermediates associated with TCA cycle, including end-stage metabolites fumarate and oxaloacetate. These data suggest STAT1 loss increases fatty acid breakdown and flux through the TCA cycle. By contrast, TCA cycle intermediates are frequently depleted (red) in the iWAT of mice fed HFD relative to normal chow controls (Figure 4F). Thus, gene expression and metabolite profiles establish *STAT1* deletion in the iWAT powers an integrated program that restores TCA cycle flux (green stars; Figure 4G) to enhance adipocyte respiration and expandability in the face of nutrient stress.

### Ablation of IFN_γ_ signaling restores insulin sensitivity and metabolic homeostasis in obese mice

The IFN_γ_gR1 protein imparts the IFN_γ_ signal to the transcription of unique proinflammatory genes. An important question is whether signals upstream of STAT1 oppose critical responses that couple WAT expandability to insulin sensitivity. To address this question, we placed *IFN_γ_gR1^−/-^* and *IFN_γ_gR1+/+* mice on HFD for 12 weeks followed by comprehensive metabolic phenotyping and mechanistic studies. Body weight (Figure 5A) and composition (Figure 5B) studies demonstrated *IFN_γ_gR1^−/-^* mice were resistant to diet-induced obesity. Accordingly, *IFN_γ_gR1* deletion improved insulin sensitivity (Figure 5C) and reduced fasting insulin levels (Figure 5D), HOMA-IR (Figure 5E). Lastly, fasting leptin levels reflected reduced fat mass in *IFN_γ_gR1^−/-^* mice (Figure 5F). Histological sections of iWAT (Figure 5G) indicated *IFN_γ_gR1^−/-^* allowed adipocyte hyperplasia with reduced cell size (Figure 5H). Consistent with the experiments in obese *STAT1^fKO^* mice (Figure 3–4), we observed suppressed IFN_γ_-STAT1 pro-inflammatory target genes coupled with enhanced expression of lipid metabolism and mitochondrial genes in iWAT, suggesting enhanced metabolic function (Figure 5I).

**Figure 5.**
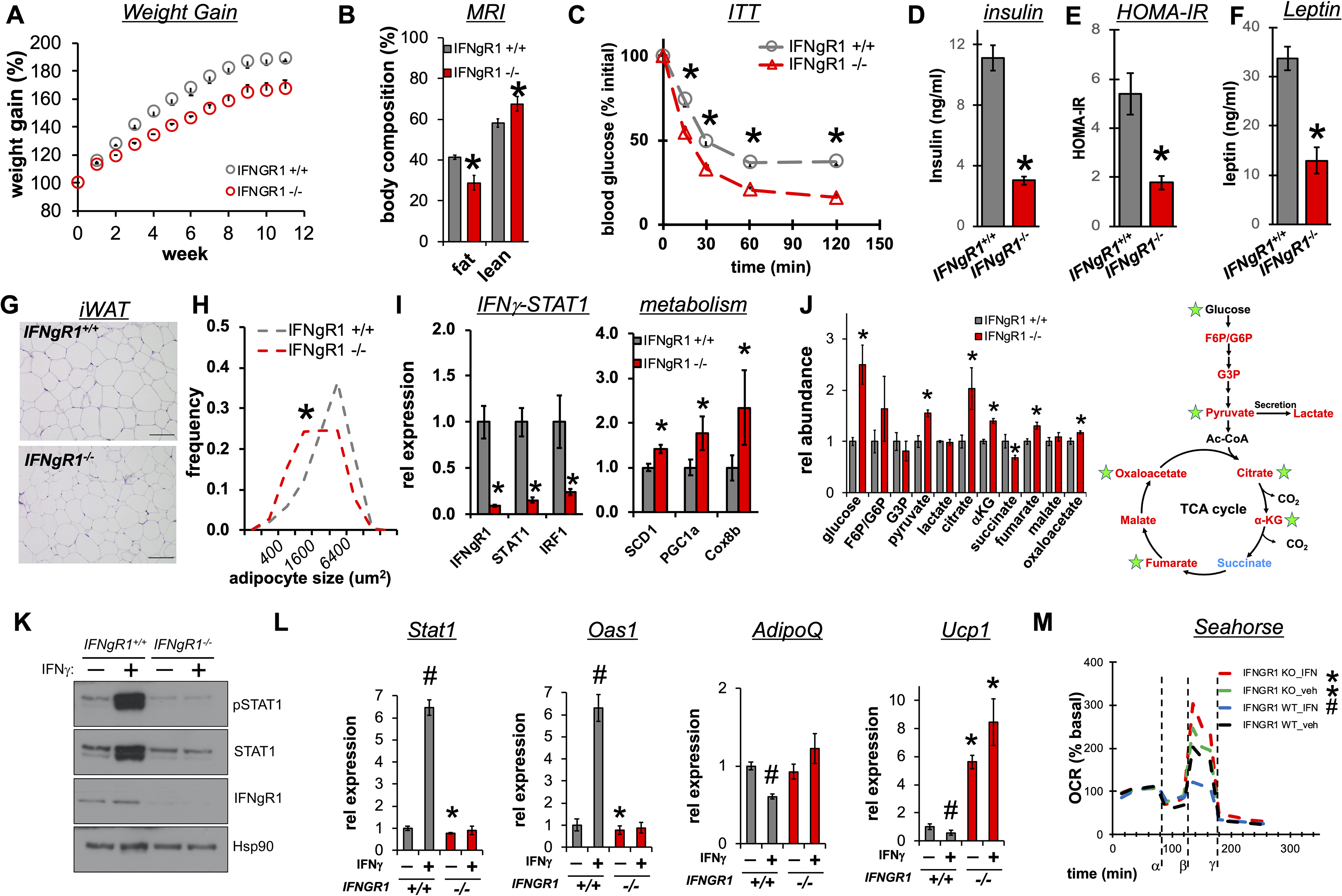
Complete disruption of IFN_γ_ signaling restores metabolic homeostasis in adipocytes and insulin sensitivity in DIO mice. **(A)** Body weight gain (% initial; n=4-5/group) and **(B)** body composition (MRI; n=9/group) measured after 12 weeks of HFD. Insulin sensitivity was determined by **(C)** insulin tolerance tests in obese *IFN_γ_gR1+/+* and *IFN_γ_gR1^−/-^* mice (n=9 mice/group, *p<0.05). **(D)** Serum insulin levels and **(E)** HOMA-IR were assessed in fasted mice (n=9 mice/group, *p<0.05). **(F)** Serum leptin levels were assessed in obese mice fasted 4 h (n=9 mice/group *p<0.05). **(G)** iWAT H/E and **(H)** adipocyte cell size distribution (% total cells) tabulated across four 20x fields of view per mouse fat depot (n=4-5/group) grey – *IFN_γ_gR1+/+*; red – *IFN_γ_gR1^−/-^*. **(I)** IFN_γ_-STAT1 inflammation and metabolism genes from iWAT of *IFN_γ_gR1+/+* and *IFN_γ_gR1^−/-^* mice on HFD (n=13-14/group). **(J)** Metabolite levels in iWAT of obese *IFN_γ_gR1+/+* (grey) and *IFN_γ_gR1^−/-^* (red) mice (n=4-5/group) were assessed using mass spectrometry. *p<0.05, data are represented as mean +/− SEM. Red metabolites decreased by HFD in WT mice; *IFN_γ_gR* deletion rescued (green) and increased (blue) metabolites. **(K)** Validation of IFN_γ_gR1 deletion and impaired STAT1 signaling by Western blot analysis of total cell lysates from *IFN_γ_gR1+/+* and *IFN_γ_gR1^−/-^* SVF-derived adipocytes after 24 h exposure to IFN_γ_. Relative mRNA expression of **(L)** *Stat1*, *Oas1*, *AdipoQ*, *Ucp1* from *IFN_γ_gR1+/+* (grey) and *IFN_γ_gR1^−/-^* (red) adipocytes after exposure to IFN_γ_ (n=3, *p<0.05 vs *IFN_γ_gR1+/+*, #p<0.05 vs vehicle). **(M)** Respiration (as oxygen consumption rate, OCR) was measured in *IFN_γ_gR1+/+* and *IFN_γ_gR1^−/-^* adipocytes after IFN_γ_ treatment and during oligomycin (α), Carbonyl cyanide-*4*-(trifluoromethoxy)phenylhydrazone (FCCP) (β), and antimycin-A/Rotenone (γ) (n=5, *p<0.05 vs *IFN_γ_gR1+/+*, #p<0.05 vs vehicle) additions. (black – *IFN_γ_gR1+/+* vehicle, blue – *IFN_γ_gR1+/+* +IFN_γ_, green – *IFN_γ_gR1^−/-^* vehicle, red – *IFN_γ_gR1^−/-^* +IFN_γ_). Data are represented as mean +/− SEM.

Disruption of IFN_γ_ activity improves mitochondrial function in adipocytes ^15,29,38^. Therefore, we performed metabolomics to examine the impact of complete ablation of IFN_γ_ signaling on the steady state levels of glycolytic and TCA cycle metabolites in iWAT. While obese *STAT1^fKO^* mice exhibited partial restoration of TCA metabolite pools (Figure 4G), *IFN_γ_gR1* deletion rescued most TCA cycle metabolites that were sensitive to a HFD, including oxaloacetate, citrate, α-ketoglutarate, and fumarate (Figure 5J). Next, to test whether *IFN_γ_gR1* deletion enabled metabolic resistance to IFN_γ_ in a cell autonomous manner, we treated differentiated SVF derived from the iWAT of either *IFN_γ_gR1−*/− mice or control mice with IFN_γ_. *IFN_γ_gR1−*/− adipocytes did not respond to IFN_γ_ at the level of p-STAT1 (Figure 5K) or canonical STAT1 target genes (Figure 5L). Accordingly, the ability of IFN_γ_ to inhibit the expression of adipocyte marker (*AdipoQ)* and mitochondrial metabolism *(Ucp1)* genes was lost in cells lacking *IFN_γ_gR1*. Metabolic activity of wild-type adipocytes remained IFN_γ_ sensitive as demonstrated by reductions in maximal oxygen consumption rate (OCR) (Figure 5M). Consistent with metabolic resistance, IFN_γ_ did not affect maximal OCR in *IFN_γ_GR1−*/− adipocytes. In summary, ablation of IFN_γ_ signaling reduces weight gain while preserving insulin sensitivity and promoting physiologic adipocyte hyperplasia. At the molecular level, ablation of IFN_γ_ signaling reduces WAT inflammation, increases levels of glycolytic and TCA metabolites, and improves respiratory capacity.

## DISCUSSION

Our work sheds light on the enigmatic role of inflammation in obesity. Numerous studies implicate cytokines and soluble mediators in the response of WAT to overnutrition and the metabolic dysfunction that characterizes obesity in rodents and humans. In this study, we demonstrate depletion of STAT1 in adipocytes reduces inflammation in WAT while enabling metabolic adaptations that promote healthy adipocyte hyperplasia. Our observations linking *STAT1* expression in subcutaneous WAT to elevated plasma glucose in humans suggests IFN_γ_ signaling influences insulin sensitivity. However, adipocyte specific STAT1 knockout exerts a nominal influence on insulin sensitivity in obese mice. Rather, we found complete disruption of IFN_γ_ signaling exerts anti-diabetic effects and enables adipocytes to retain metabolic function in obesity and T2DM.

Obesity and insulin resistance increase IFN_γ_ production ^39^ and IFN_γ_-mediated activation of the transcription factor STAT1 blocks adipocyte differentiation ^15,21,29,38^. We extend these findings with genetic and RNA interference studies *in vitro* that demonstrate STAT1 depletion increases human and mouse adipocyte differentiation. To our surprise, STAT1 deletion in WAT marginally improved the metabolic criteria associated with obesity. Although STAT1 mediates many IFN_γ_-dependent actions, IFN_γ_ also alters expression of many genes in STAT1^−/-^ cells ^38,40^. Indeed, complete elimination of IFN_γ_ activity improves insulin sensitivity in obese mice. Although whole body *IFN_γ_gR1* deletion exposes impacts on immunity across multiple endocrine tissues ^17,39,41^, *IFN_γ_gR1−*/− adipocytes resist the detrimental effects of IFN_γ_ on metabolism and nominate cell autonomous functions that restrict WAT expandability in obesity.

Inflammation and decreased mitochondrial oxidative capacity frequently co-present in obesity. Interferons broadly inhibit expression of several mitochondrial genes encoded within the mitochondrion ^18–20,29^. The mechanisms by which interferon activation reduces mitochondrial function remain unclear. One possibility is that IFN_γ_-activated STAT1 represses the transcription of integral regulators of lipid and glucose metabolism, as occurs with PPAR_γ_, in obesity. IFN_γ_ causes STAT1 binding near PPAR_γ_ sites in multiple enhancer regions of mitochondrial and insulin sensitivity genes corresponding with reduced mRNA expression ^15,42^. These findings suggest a negative crosstalk between the occupancy of PPAR_γ_ and STAT1 binding sites near genes important for WAT expansion and whole-body insulin sensitivity.

Reduced subcutaneous adipocyte differentiation in hypertrophic obesity reflects persistent mitochondrial dysfunction and diminished lipid storage ^43^. Our study and others ^44–46^ highlight how obesity slows TCA cycle function and likely impacts anabolic functions of WAT. The TCA cycle provides intermediates and energy that drive lipid synthesis in adipose tissue. To this end, failure of integral TCA and lipid metabolism reactions reflect low subcutaneous adipocyte differentiation and thus limits WAT expandability in the face of obesity. Previous animal and human studies corroborate critical effects of IFN_γ_ on endocrine tissues ^16,41,47^. The importance of this cytokine in host defense and its influence on fuel mobilization reaffirm the link between inflammatory response and energy balance disruption. Interferons lower cellular capacity to generate ATP ^18–20,29^ mirrored by decreased mitochondrial ETC activity. As a result, adipocytes cannot devote necessary ATP to biosynthesis of lipid and mitochondrial building blocks required for adipocyte expansion in response to chronic nutrient excess. Elimination of IFN_γ_ action in adipocytes repletes critical TCA intermediates. For example, accumulation of -ketoglutarate and fumarate allows generation of reducing equivalents to transfer electrons to the mitochondrial respiratory chain for subsequent production of ATP. The elevation of citrate supplies six-carbon backbones to re-establish cytosolic acetyl-CoA and oxaloacetate pools to promote lipid and nucleotide synthesis. The consequence of these events is evidenced functionally by enrichment of oxidative phosphorylation genes, superior oxygen consumption rates *in vitro*, and adipocyte differentiation in hyperplastic obesity.

In summary, these findings describe a mechanism that allows insulin resistance to occur independent of WAT inflammation. Pro-inflammatory cytokines inhibit insulin signaling and mitochondrial function in adipocytes. Also, numerous studies established strong correlations between pro-inflammatory cytokines, like TNF_α_ ^48,49^ and IFN_γ_ ^39^ and energy balance defects in adipose tissue. However, TNF_α_ inhibitors and other or broad anti-inflammatory strategies that impact STAT1 or STAT3 activity lack clinical efficacy in the treatment of obesity ^50–53^. Furthermore, recent provocative studies argue WAT inflammation responds to insulin resistance ^54^. Our studies and others ^54^ argue the direct, causative relationship between chronic WAT inflammation and insulin resistance remains oversimplified. Although our findings do not rule out the idea that inflammation degrades metabolic fitness and insulin resistance, they bring into question whether anti-inflammatory monotherapies in WAT will be an effective strategy to improve all the clinical phenotypes of obesity and T2DM.

## Author Contributions

A.R.C. and S.M.H. conceptualized the study. P.M., N.C., A.R.C., D.A.B., and S.M.H. designed experiments. A.R.C., D.A.B., and S.M.H. wrote the manuscript with editorial input from all authors. S.M.H. and A.R.C performed all experiments with assistance as noted: P.K.S. assisted with mouse phenotyping, V.P. and N.P. assisted with metabolomics analysis, N.C. performed qPCR and gene expression validation, K.K. performed analysis of liver lipid and histology, K.R. and C.C. assisted with RNA-Seq data analysis and metabolomics data integration, D.V. and R.V. provided clinical specimens. All work was performed under the supervision of S.M.H.

## REFERENCES

1. Afshin, A., et al. Health effects of overweight and obesity in 195 countries over 25 years. N Engl J Med 377, 13–27 (2017).

2. Smith, G.I., Mittendorfer, B. & Klein, S. Metabolically healthy obesity: facts and fantasies. J Clin Invest 129, 3978–3989 (2019).

3. Patsouris, D., et al. Ablation of CD11c-positive cells normalizes insulin sensitivity in obese insulin resistant animals. Cell Metab 8, 301–309 (2008).

4. Lee, Y.S., et al. Inflammation is necessary for long-term but not short-term high-fat diet-induced insulin resistance. Diabetes 60, 2474–2483 (2011).

5. Kanda, H., et al. MCP-1 contributes to macrophage infiltration into adipose tissue, insulin resistance, and hepatic steatosis in obesity. J Clin Invest 116, 1494–1505 (2006).

6. Weisberg, S.P., et al. Obesity is associated with macrophage accumulation in adipose tissue. J Clin Invest 112, 1796–1808 (2003).

7. Ouchi, N., Parker, J.L., Lugus, J.J. & Walsh, K. Adipokines in inflammation and metabolic disease. Nat Rev Immunol 11, 85–97 (2011).

8. Crewe, C., An, Y.A. & Scherer, P.E. The ominous triad of adipose tissue dysfunction: inflammation, fibrosis, and impaired angiogenesis. J Clin Invest 127, 74–82 (2017).

9. Kintscher, U., et al. T-lymphocyte infiltration in visceral adipose tissue: a primary event in adipose tissue inflammation and the development of obesity-mediated insulin resistance. Arterioscler Thromb Vasc Biol 28, 1304–1310 (2008).

10. Wentworth, J.M., et al. Interferon-gamma released from omental adipose tissue of insulin-resistant humans alters adipocyte phenotype and impairs response to insulin and adiponectin release. Int J Obes 41, 1782–1789 (2017).

11. Majoros, A., et al. Response to interferons and antibacterial innate immunity in the absence of tyrosine-phosphorylated STAT1. EMBO Rep 17, 367–382 (2016).

12. Qing, Y. & Stark, G.R. Alternative activation of STAT1 and STAT3 in response to interferon-gamma. J Biol Chem 279, 41679–41685 (2004).

13. Wong, L.H., et al. Isolation and characterization of a human STAT1 gene regulatory element. Inducibility by interferon (IFN) types I and II and role of IFN regulatory factor-1. J Biol Chem 277, 19408–19417 (2002).

14. Cheon, H. & Stark, G.R. Unphosphorylated STAT1 prolongs the expression of interferon-induced immune regulatory genes. Proc Natl Acad Sci U S A 106, 9373–9378 (2009).

15. Koh, E.H., et al. miR-30a remodels subcutaneous adipose tissue inflammation to improve insulin sensitivity in obesity. Diabetes 67, 2541–2553 (2018).

16. O’Rourke, R.W., et al. Systemic inflammation and insulin sensitivity in obese IFN-gamma knockout mice. Metabolism 61, 1152–1161 (2012).

17. Deng, T., et al. Adipocyte adaptive immunity mediates diet-induced adipose inflammation and insulin resistance by decreasing adipose Treg cells | Nature Communications. Nat Commun 8, 15725 (2017).

18. Hahn, W.S., et al. Proinflammatory cytokines differentially regulate adipocyte mitochondrial metabolism, oxidative stress, and dynamics. Am J Physiol Endocrinol Metab 306, E1033–1045 (2014).

19. Kissig, M., et al. PRDM16 represses the type I interferon response in adipocytes to promote mitochondrial and thermogenic programing. EMBO J 36, 1528–1542 (2017).

20. Lewis, J.A., Huq, A. & Najarro, P. Inhibition of mitochondrial function by interferon. J Biol Chem 271, 13184–13190 (1996).

21. McGillicuddy, F.C., et al. Interferon gamma attenuates insulin signaling, lipid storage, and differentiation in human adipocytes via activation of the JAK/STAT pathway. J Biol Chem 284, 31936–31944 (2009).

22. Klover, P.J., et al. Loss of STAT1 from mouse mammary epithelium results in an increased Neu-induced tumor burden. Neoplasia 12, 899–905 (2010).

23. Colleluori, G., et al. Adipocytes ESR1 expression, body fat and response to testosterone therapy in hypogonadal men vary according to estradiol devels. Nutrients 10, pii: E1226 (2018).

24. Galarraga, M., et al. Adiposoft: automated software for the analysis of white adipose tissue cellularity in histological sections. J Lipid Res 53, 2791–2796 (2012).

25. Trapnell, C., et al. Transcript assembly and quantification by RNA-Seq reveals unannotated transcripts and isoform switching during cell differentiation. Nat Biotechnol 28, 511–515 (2010).

26. Kim, D., Langmead, B. & Salzberg, S.L. HISAT: a fast spliced aligner with low memory requirements. Nat Methods 12, 357–360 (2015).

27. Koh, E.H., et al. Mitochondrial Activity in Human White Adipocytes Is Regulated by the Ubiquitin Carrier Protein 9/microRNA-30a Axis. J Biol Chem 291, 24747–24755 (2016).

28. Bader, D.A., et al. Mitochondrial pyruvate import is a metabolic vulnerability in androgen receptor-driven prostate cancer. Nat Metab 1, 70–85 (2019).

29. Moisan, A., et al. White-to-brown metabolic conversion of human adipocytes by JAK inhibition. Nat Cell Biol 17, 57–67 (2015).

30. Dahlman, I., et al. Downregulation of electron transport chain genes in visceral adipose tissue in type 2 diabetes independent of obesity and possibly involving tumor necrosis factor-alpha. Diabetes 55, 1792–1799 (2006).

31. Grohmann, M., et al. Obesity drives STAT-1-dependent NASH and STAT-3-dependent HCC. Cell 175, 1289–1306.e1220 (2018).

32. Chen, T.T., et al. STAT1 regulates marginal zone B cell differentiation in response to inflammation and infection with blood-borne bacteria. J Exp Med 213, 3025–3039 (2016).

33. Johnson, L.M. & Scott, P. STAT1 expression in dendritic cells, but not T cells, is required for immunity to Leishmania major. J Immunol 178, 7259–7266 (2007).

34. Kang, Y.H., Biswas, A., Field, M. & Snapper, S.B. STAT1 signaling shields T cells from NK cell-mediated cytotoxicity. Nat Commun 10, 912 (2019).

35. Eguchi, J., et al. Transcriptional control of adipose lipid handling by IRF4. Cell Metab 13, 249–259 (2011).

36. Cox, A.R., Chernis, N., Masschelin, P.M. & Hartig, S.M. Immune Cells Gate White Adipose Tissue Expansion. Endocrinology 160, 1645–1658 (2019).

37. Koves, T.R., et al. Mitochondrial overload and incomplete fatty acid oxidation contribute to skeletal muscle insulin resistance. Cell Metab 7, 45–56 (2008).

38. Todoric, J., et al. Cross-talk between interferon-gamma and hedgehog signaling regulates adipogenesis. Diabetes 60, 1668–1676 (2011).

39. Reardon, C.A., et al. Obesity and insulin resistance promote atherosclerosis through an IFN gamma-regulated macrophage protein network. Cell Rep 23, 3021–3030 (2018).

40. Gil, M.P., et al. Biologic consequences of Stat1-independent IFN signaling. Proc Natl Acad Sci U S A 98, 6680–6685 (2001).

41. Rocha, V.Z., et al. Interferon-gamma, a Th1 cytokine, regulates fat inflammation: a role for adaptive immunity in obesity. Circ Res 103, 467–476 (2008).

42. Siersbaek, R., et al. Transcription factor cooperativity in early adipogenic hotspots and super-enhancers. Cell Rep 7, 1443–1455 (2014).

43. Hammarstedt, A., Gogg, S., Hedjazifar, S., Nerstedt, A. & Smith, U. Impaired adipogenesis and dysfunctional adipose tissue in human hypertrophic obesity. Physiol Rev 98, 1911–1941 (2018).

44. Cummins, T.D., et al. Metabolic remodeling of white adipose tissue in obesity. Am J Physiol Endocrinol Metab 307, E262–277 (2014).

45. Petrus, P., et al. Glutamine links obesity to inflammation in human white adipose tissue. Cell Metab 31, 375–390 (2020).

46. Neeland, I.J., et al. Metabolomics profiling of visceral adipose tissue: Results from MESA and the NEO study. J Am Heart Assoc 8, e010810 (2019).

47. Memon, R.A., Feingold, K.R., Moser, A.H., Doerrler, W. & Grunfeld, C. In vivo effects of interferon-alpha and interferon-gamma on lipolysis and ketogenesis. Endocrinology 131, 1695–1702 (1992).

48. Hotamisligil, G.S., Arner, P., Caro, J.F., Atkinson, R.L. & Spiegelman, B.M. Increased adipose tissue expression of tumor necrosis factor-alpha in human obesity and insulin resistance. J Clin Invest 95, 2409–2415 (1995).

49. Hotamisligil, G.S., Shargill, N.S. & Spiegelman, B.M. Adipose expression of tumor necrosis factor-alpha: direct role in obesity-linked insulin resistance. Science 259, 87–91 (1993).

50. Goldfine, A.B., et al. The effects of salsalate on glycemic control in patients with type 2 diabetes: a randomized trial. Ann Intern Med 152, 346–357 (2010).

51. Ofei, F., Hurel, S., Newkirk, J., Sopwith, M. & Taylor, R. Effects of an engineered human anti-TNF-alpha antibody (CDP571) on insulin sensitivity and glycemic control in patients with NIDDM. Diabetes 45, 881–885 (1996).

52. Goldfine, A.B., et al. A randomised trial of salsalate for insulin resistance and cardiovascular risk factors in persons with abnormal glucose tolerance. Diabetologia 56, 714–723 (2013).

53. Wascher, T.C., et al. Chronic TNF-alpha neutralization does not improve insulin resistance or endothelial function in “healthy” men with metabolic syndrome. Mol Med 17, 189–193 (2011).

54. Shimobayashi, M., et al. Insulin resistance causes inflammation in adipose tissue. J Clin Invest 128, 1538–1550 (2018).

